# Identification of within-host deletions in domain 0 of the spike gene of highly pathogenic feline coronavirus type 2 from the United States

**DOI:** 10.1101/2025.07.12.664542

**Authors:** Ximena A. Olarte-Castillo, Abigail Schlecht, Kelly Sams, Laura B. Goodman, Gary R. Whittaker

## Abstract

Feline coronavirus (FCoV) is known to gain pathogenicity within-host to cause the lethal disease feline infectious peritonitis (FIP). Most FIP cases are caused by viruses in genotype 1 (FCoV-1) via an ‘internal mutation’ in the spike gene. However, genotype 2 (FCoV-2) has risen to prominence based on the emergence of FCoV-23, a highly pathogenic novel variant from Cyprus that has a deletion in the N-terminus (domain 0) of spike. Here, we conducted a retrospective molecular study of FCoV-2 detected in three cats in the U.S. during 2013 and 2016. Whole-genome sequencing revealed that the two cats exhibiting long-term signs each had an FCoV-2 with a distinct deletion in domain 0 of spike in all examined tissues. The epidemiologically-linked cat displaying signs for a short duration had an FCoV-2 with an intact spike. Our results suggest that this “internal deletion” in the spike gene is a biomarker of highly pathogenic FCoV-2.

**Graphical abstract:** 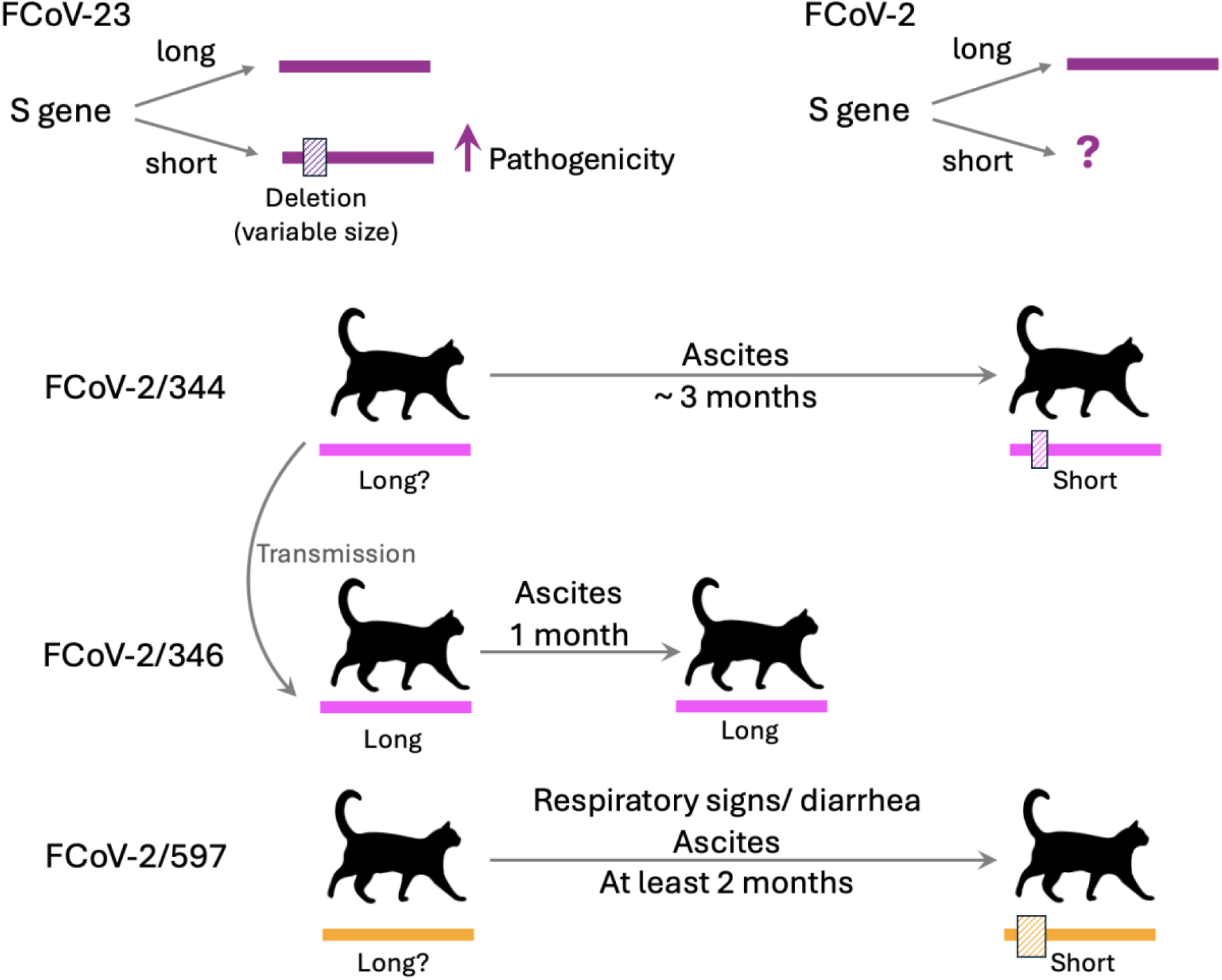

## Introduction

Feline and canine coronavirus (FCoV and CCoV, respectively) are alphacoronaviruses highly prevalent in domestic cat and dog populations worldwide. FCoV and CCoV are classified into two genotypes, types 1 and 2, with extensive recombination occurring between them (1). For example, FCoV type 2 (FCoV-2) is a genotype resulting from the recombination between FCoV type 1 (FCoV-1) and CCoV type 2 (CCoV-2 (2)). Both FCoV-1 and FCoV-2 are found in two biotypes: a generally apathogenic feline enteric coronavirus (FECV) and a highly pathogenic, macrophage-tropic form known as feline infectious peritonitis virus (FIPV (3)). Infection with the highly pathogenic form leads to feline infectious peritonitis (FIP), a disease that is usually fatal without antiviral treatment (4). FCoV-1 is more widespread among domestic cats worldwide, whereas various recombinant variants of FCoV-2 have mostly been reported in small outbreaks (5, 6). While FCoV infection is common in multi-cat settings, affecting nearly 80% of cats, only a small fraction (about 5%) will develop FIP (7). In 2023, a large-scale outbreak of a novel FCoV-2-like recombinant variant (known as FCoV-23) swept through the free-ranging cat population in the Republic of Cyprus (8). The prevalence remains unclear, with estimates ranging from 10,000 (8) to as many as 300,000 (9) cases of FIP. Isolated cases of FCoV-23 have been introduced into the UK through travel (10), and the spread of this highly pathogenic FCoV poses a threat to global cat populations, warranting surveillance of this and similar emerging viruses in other countries. The spike (S) protein of coronaviruses plays a crucial role in pathogenicity by facilitating host cell entry through receptor binding (S1 domain) and membrane fusion (S2 domain). For FCoV-1, pathogenicity has been linked to mutations generated within individual cats (11), with key mutation(s) now thought to lie within the S gene (12). FCoV-1 has a cleavage site between S1 and S2 (the S1/S2 cleavage site), which primes the virus for fusion (13). In transmissible, low-pathogenicity FCoVs, this cleavage site is characterized by a consensus motif (SRRSRR|S), where R is arginine, S is Serine, and | indicates the cleavage site. Point mutations disrupting this motif and reducing the proteolytic priming of S are strongly linked to pathogenicity (12). Other markers of pathogenicity for FCoV-1 include a mutation in a residue located in the S2 domain (known as M1058L (14)) and at the S2’ cleavage site (15), although the role of these mutations remains unclear. The “internal mutation” hypothesis suggests that the accumulation of these (and possibly other) mutations enables the virus to gain macrophage tropism (11). FCoV-2 and FCoV-23 lack the S1/S2 cleavage site and the ‘1058’ residue, and there is less understanding of the molecular mechanisms underlying the emergence of pathogenic variants (16, 17).

Two characteristics define the S of FCoV-23: (1) it likely originated from a ‘pantropic’ CCoV-2 (pCCoV-2) known to circulate in the Mediterranean basin, with pathogenic CCoV-2 NA-09 detected in Greece in 2009 being its closest relative (18), and (2) two versions of spike have been identified: a long one and a short one (8). The short version has a deletion in the N-terminus of the S protein that encompasses most of the domain 0. This deletion has been reported in about 90% of FCoV-23/FIP cats and varies in size among individuals (ranging from 152 to 245 amino acids), indicating that each deletion develops independently within each cat (8). To date, these deletions have not been reported in other FCoV-2 (though only seven complete FCoV-2 S sequences are currently available, Appendix Table 1). The deletion contributes to the pathogenicity of this novel FCoV by enhancing fusogenicity and accelerating viral cell entry in feline and canine tissues (19).

A distinctive feature of Cyprus is its large population of free-roaming cats, estimated to be up to 2 million, which may be one of the factors contributing to the rapid spread of FCoV-23 (8). However, large populations of free-roaming domestic cats also exist in many cities across the U.S., including New York City (estimated at up to 1 million (20)) and Washington D.C. (estimated at around 200,000), where a program to count the free-ranging cat population has been recently established (21). Thus, identifying and understanding the potential impact of FCoV-2 and FCoV-2-like viruses circulating in the U.S. is crucial for understanding the risk of emergence and spreading of pathogenic variants.

In this study, we conducted a retrospective analysis of three cases of FCoV-2 in the U.S.: two cases (#344 and #346) from 2013 and one (#597) from 2016. Cats #344 and #597 exhibited clinical signs consistent with FCoV infection for at least 2 months. The third cat (#346) was linked epidemiologically to cat #344 and was euthanized shortly after showing signs. Whole genome sequencing revealed that the cats showing long-term signs (#344 and #597) each had an FCoV-2 with a distinct deletion in S, which was found in all examined tissues. In contrast, cat #346 had an FCoV-2 with the long version of S. Phylogenetic analysis indicated cats #344 and #346 were infected with the same FCoV-2. The only difference was that the FCoV-2 detected in #346 lacked the deletion S. Although the FCoV-2 with the short version was found in the feces of cats #344 and #597, its RNA was found in very low loads (2.52 to 5.28 RNA copies/μl). Our data are consistent with a model whereby FCoV-2 displaying the long version of S is transmitted between cats, while the short version is generated within each cat after a prolonged infection and spreads rapidly throughout the body. We refer to this as the “internal deletion” hypothesis. This study is the first report of FCoV-2 displaying short versions of S in the U.S. and indicates that this mechanism is not exclusive to the evolution of FCoV-23.

## Materials and Methods

### Sample collection and processing

Tissue, feces, and ascites were collected in 2013 from cats #344 and #346 in Plainview, Arkansas, and in 2016 from cat #597 in Philadelphia, Pennsylvania. Cat #344 was a three-year-old British short-haired (BSH) female residing in a multi-cat environment with five other cats. In early March 2013, she started showing signs of ascites and was euthanized on 23 May 2013. Cat #346 was a 9-month-old BSH female and the daughter of #344, who started showing signs of ascites on 6 May 2013, and was euthanized on 4 June 2013. Cat #597 was an 11-month-old short-haired male residing in a multi-cat environment with 12 other cats. The cat had a history of upper respiratory infections and diarrhea. On 15 March 2016, he started showing signs of ascites and was euthanized on 5 May 2016. Histological tissue processing and detection of FCoV by immunohistochemistry were done at the Cornell University College of Veterinary Medicine Animal Health Diagnostic Center. After collection, samples were frozen at -80°C until further use.

### FCoV screening, RNA quantification, and whole genome sequencing

RNA was extracted from the 20 collected samples (Table 1), as described previously (22, 23). FCoV RNA was detected using the TaqMan Fast Virus 1-Step Master Mix for qPCR (Thermo Fisher Scientific) and previously published primers/probe (24). This assay targets the intergenic region between the matrix and nucleocapsid genes and does not differentiate between FCoV-1 and FCoV-2. Whole genomes of FCoV were sequenced using hybridization capture with a panel of 3,140 baits targeting FCoV-1, FCoV-2, CCoV-1, and CCoV-2 (design ID: TE-92694309, Twist Biosciences). The synthesis of dsDNA, Illumina-compatible libraries, hybridization capture, and next-generation sequencing were performed as previously described (22). Paired-end reads were trimmed for quality using Trimmomatic (25). Trimmed reads were mapped against 115 sequences of FCoV-1, FCoV-2, FCoV-23, CCoV-2, CCoV-1, and TGEV (Appendix Table 2) using BWA-MEM2 (26). Quality trimming and mapping were done in Galaxy V22.01 (27). Viral RNA was quantified using the QX200 droplet digital PCR (ddPCR) System, the 1-Step RT-ddPCR Advanced Kit for Probes (Biorad), and the primers and probes shown in Appendix Table 3.

**Table 1.**
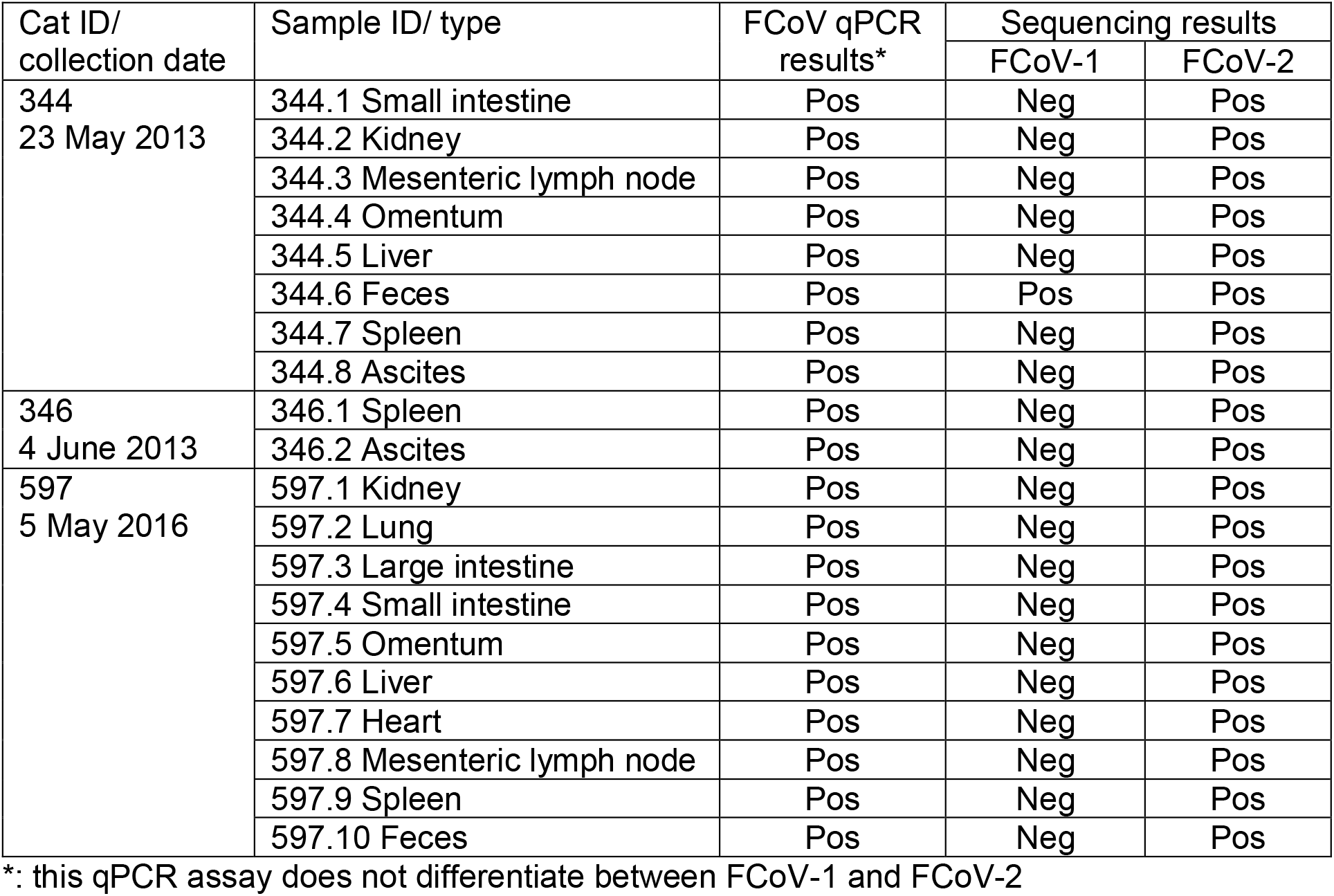
Samples collected from cats #344, #346, and #597. The results from the screening and sequencing of FCoV for each sample are also shown.

### Genetic analyses

The 21 whole genome sequences obtained in this study were aligned with 115 additional whole genomes of selected FCoV-1, FCoV-2, FCoV-23, CCoV-1, CCoV-2, and TGEV (Appendix Table 2) using the MUSCLE algorithm (28). Percentage similarity between sequences was calculated in Geneious Prime ® 2023.2.1 (Biomatters Ltd), recombination breakpoints were detected in the Recombination Detection Program (RDP) 5 (29) using the RDP method (30), and similarity plots were generated in SimPlot 5.1 (31). The complete ORF1a region (11,613 nt) was aligned for the three sequences detected in this study and 37 others from FCoV-23, FCoV-2, FCoV-1, and CCoV-2 (Appendix Table 3). This alignment was used to construct a maximum likelihood (ML) phylogeny using Mega 12.0.11 (32). The amino acid sequence of the complete S protein was aligned for the three sequences obtained in this study and 40 others of FCoV-23, FCoV-2, and CCoV-2 (Appendix Table 2) using the Clustal Omega algorithm (33). Excluding domain 0, the alignment of the S protein was used to construct an ML phylogeny using Mega 12.0.11 (32). All phylogenies were visualized and color-edited using iTol v7.2 (34). Both the alignment of the complete S protein and the one excluding domain 0 were used to generate a similarity plot using SimPlot 5.1 (31).

**Table 2.** Size (in amino acids) of the 11 open reading frames (ORFs) of the detected FCoV-2 and FCoV-23. The three sequences obtained in this study are in bold.

| Variant name<br>(Accession number) | ORF1a | ORF1b | S | ORF3a | ORF3b | ORF3c | E | M | N | 7a | 7b |
| --- | --- | --- | --- | --- | --- | --- | --- | --- | --- | --- | --- |
| **FCoV-2 344 USA 2013** | **4032** | **2611** | **1295** | **105** | **43** | **67** | **82** | **291** | **376** | **108** | **206** |
| **FCoV-2 346 USA 2013** | **4025** | **2611** | **1453** | **105** | **43** | **244** | **82** | **288** | **376** | **108** | **206** |
| **FCoV-2 597 USA 2016** | **4024** | **2611** | **1240** | **70** | **73** | **208** | **82** | **291** | **377** | **108** | **206** |
| FCoV23 C8/Wi/8299 (PQ133181) | 4029 | 2611 | 1243 | 70 | 73 | 237 | 82 | 291 | 377 | 108 | 206 |
| FCoV23 D8/Ta/8364 (PQ133184) | 4029 | 2611 | 1241 | 70 | 73 | 237 | 82 | 291 | 377 | 108 | 206 |
| FCoV23 F9/Fe/9265 (PQ133190) | 4029 | 2611 | 1241 | 70 | 73 | 237 | 82 | 291 | 377 | 108 | 206 |
| FCoV23 E9/Mi/9160 (PQ133188) | 4029 | 2611 | 1240 | 70 | 73 | 237 | 82 | 291 | 377 | 108 | 206 |
| FCoV23 F11/Gi/6590 (PQ133176) | 4029 | 2611 | 1235 | 70 | 73 | 237 | 82 | 291 | 377 | 108 | 206 |
| FCoV23 D9/Ni/9183 (PQ133185) | 4029 | 2611 | 1236 | 70 | 73 | 237 | 82 | 291 | 377 | 108 | 206 |
| FCoV23 H8/So/8842 (PQ133194) | 4029 | 2611 | 1238 | 70 | 73 | 237 | 82 | 291 | 377 | 108 | 206 |
| FCoV23 A9/Do/9045 (PQ133179) | 4029 | 2611 | 1250 | 70 | 73 | 237 | 82 | 291 | 377 | 108 | 206 |
| FCoV23 C11/Re/10276 (PQ133182) | 4029 | 2611 | 1454 | 70 | 73 | 237 | 82 | 291 | 377 | 108 | 206 |
| FCoV23 F12/BW/11350 (PQ133177) | 4029 | 2611 | 1224 | 70 | 73 | 237 | 82 | 291 | 377 | 108 | 206 |

## Results

Twenty samples were collected from the three cats, and all tested positive for FCoV by qPCR (Table 1). Whole genome sequencing, recombination analyses, and similarity plots revealed that all the FCoVs detected were recombinants between FCoV-1 and CCoV-2 (i.e. FCoV-2, Figure 1). The recombinant region of the FCoV-2 detected in cats #344, and #346 (FCoV-2/344, FCoV-2/346, respectively) covered most of the ORF1b and 3abc genes (RDP, p <0.001), and the one from cat #597 (FCoV-2/597) only included the S gene (RDP, p <0.001). In the fecal sample of cat #344 we detected co-infection with FCoV-1 (FCoV-1/344.6, Table 1). The S1/S2 cleavage site of FCoV-1/344.6 was SRRSRR|S, and residue ‘1058’ was M, consistent with the shedding of a low pathogenicity phenotype. The FCoV-2s detected in all the tissues of each cat were nearly identical (>99.9% similarity); therefore, only one sequence for each FCoV-2 will be used as a reference from this point forward.

**Figure 1.**
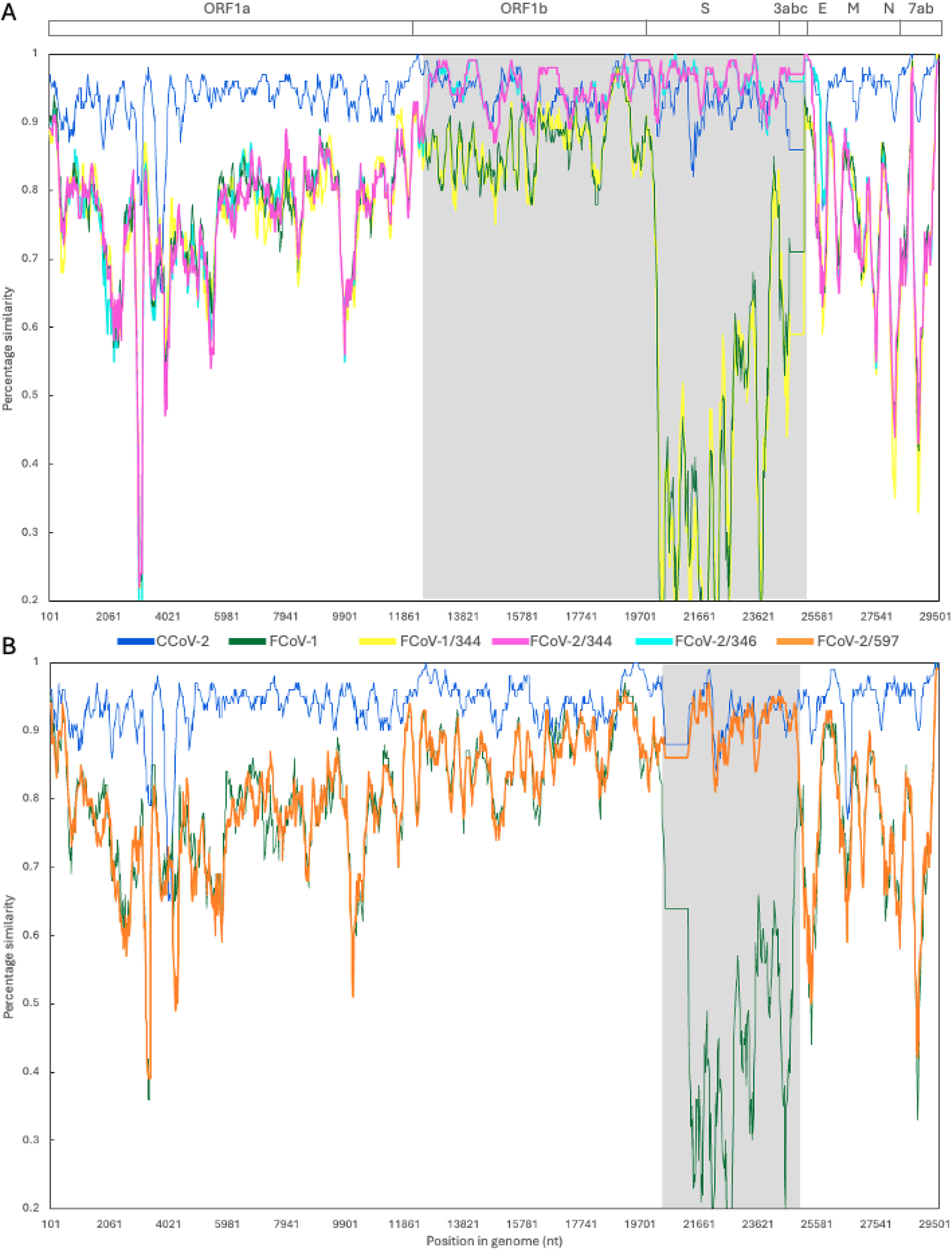
Similarity plots of the whole genome sequences of FCoV-1/344, FCoV-2/344, FCoV-2/346 (A), and (B) FCoV-2/597. On top is a graphical depiction of the general organization of the genome of FCoV/CCoV, including the non-structural (ORF1a, ORF1b), structural (S, E, M, N), and accessory (3a, b, c, and 7a, b) genes. In each graph, the nucleotide similarity across the entire genome (∼29,500 nt) is compared between a reference sequence (CCoV-2 CB-05/Italy/20025/KP981644) and the sequences obtained in this study (FCoV-1/344 in yellow, FCoV-2/344 in fuchsia, FCoV-2/346 in cyan, FCoV-2/597 in orange), FCoV-1 (in green), and CCoV-2 (in blue). Recombinant regions are shaded in gray and include the ORF1b, S, and 3abc genes for FCoV-2/344 and 346, and the S gene for FCoV-2/597. In these regions, the detected FCoV-2s are more similar to CCoV-2 (in blue) than to FCoV-1 (in green). FCoV-1/344 is more similar to FCoV-1 throughout the whole genome. The plots were generated using the Kimura 2-parameter distance model, with a 250 bp window and a 40 bp step. Sequences used in the plot: FCoV-1 RM/USA/2012/JQ404410, CCoV-2 TN449/USA/2012/JQ404410, FCoV-2 WSU79-1683/USA/1979/JN634064.

Phylogenetic analysis of ORF1a (Figure 2) showed that FCoV-1/344, FCoV-2/344, and FCoV-2/346 form a single group, suggesting a common origin for these FCoVs. FCoV-2/597 belonged to the same group as these three FCoVs (Figure 2). Genetic analysis of the S gene of the three FCoV-2 detected showed that FCoV-2/344 has a deletion of 459 nt (153 amino acids), and FCoV-2/597 has a deletion of 636 nt (212 amino acids, Figure 3). Each deletion was detected in all the tissues processed from each individual (Table 1). No deletion was detected in the S of FCoV-2/346 (Figure 3). No co-infection with the long and short versions was detected in any of the processed tissues. When compared to FCoV-23 (Table 2), major deletions were also detected in ORF3b (30 nt) and ORF3c (170 nt) of FCoV-2/344, ORF3b (30 nt) of FCoV-2/346, and ORF3c (29 nt) of FCoV-2/597 (Table 2). ORF3a of FCoV-3/344 and FCoV-2/346 was predicted to be longer (35 nt). Excluding the deletions identified in the various FCoV-2 variants, the genomes of FCoV-2/344 and FCoV-2/346 exhibited a similarity of 97.4%. Excluding the recombinant region, FCoV-1/344 and FCoV-2/344 were 94.1% similar. Excluding domain 0, the overall percentage amino acid similarity of S between FCoV-2/344, FCoV-2/597, and FCoV-23 was 97.3%. The overall percentage amino acid similarity of the complete S between FCoV-2/346 and the long version of FCoV-23 was 93.9%.

**Figure 2.**
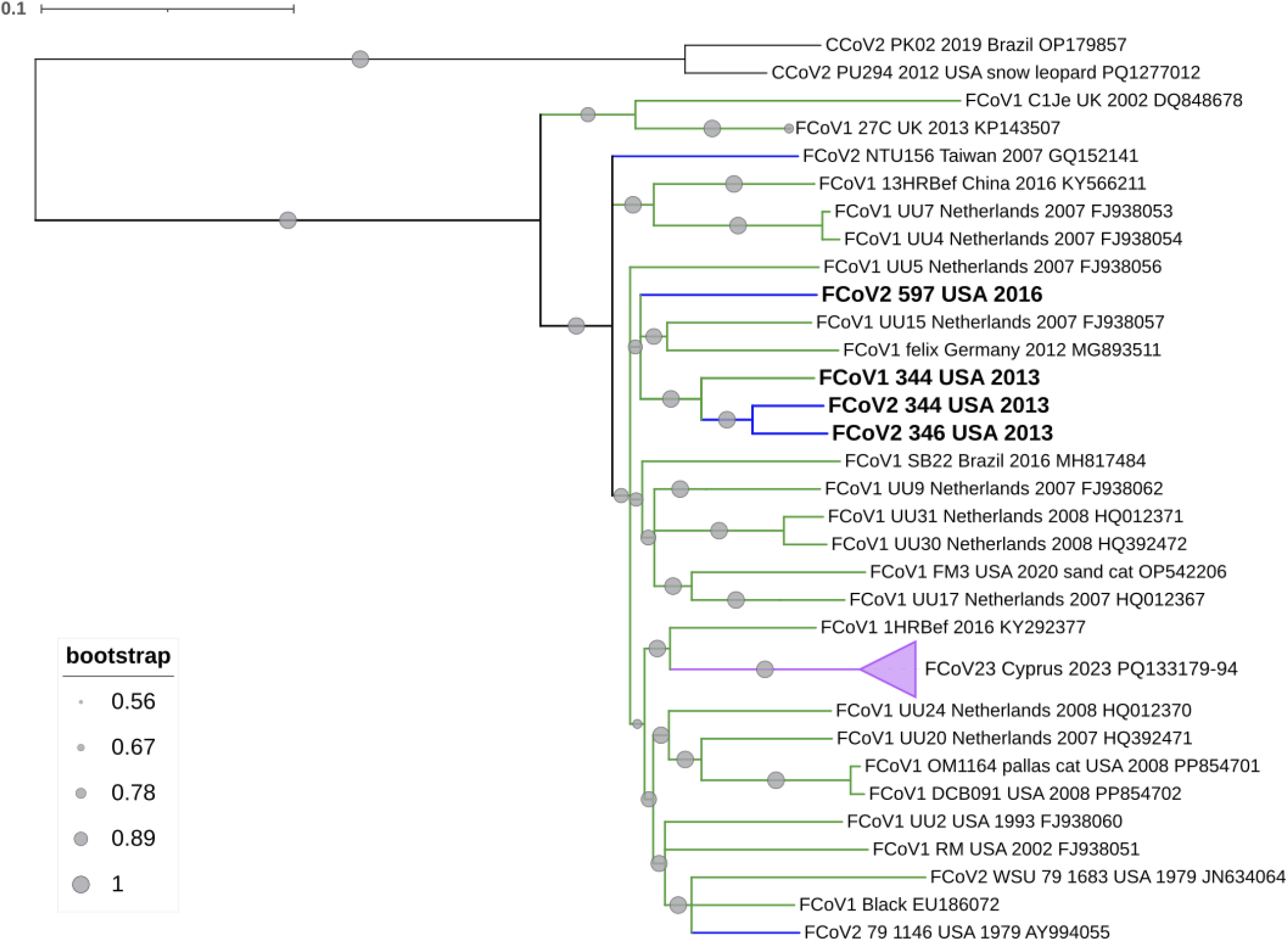
Maximum likelihood phylogenetic tree of ORF1a (11,613 nt) of FCoV and CCoV. The FCoVs sequenced in this study (344, 346, 597) are in bold. FCoV-1 is in green, FCoV-2 is in blue, and FCoV-23 is in purple. Circles in the branches indicate bootstrap percentage values for 1,000 replicates. The size of each circle is proportional to the bootstrap value, and the scale is displayed at the bottom left. Best-fitting nucleotide substitution model: GTR+G+I.

**Figure 3.**
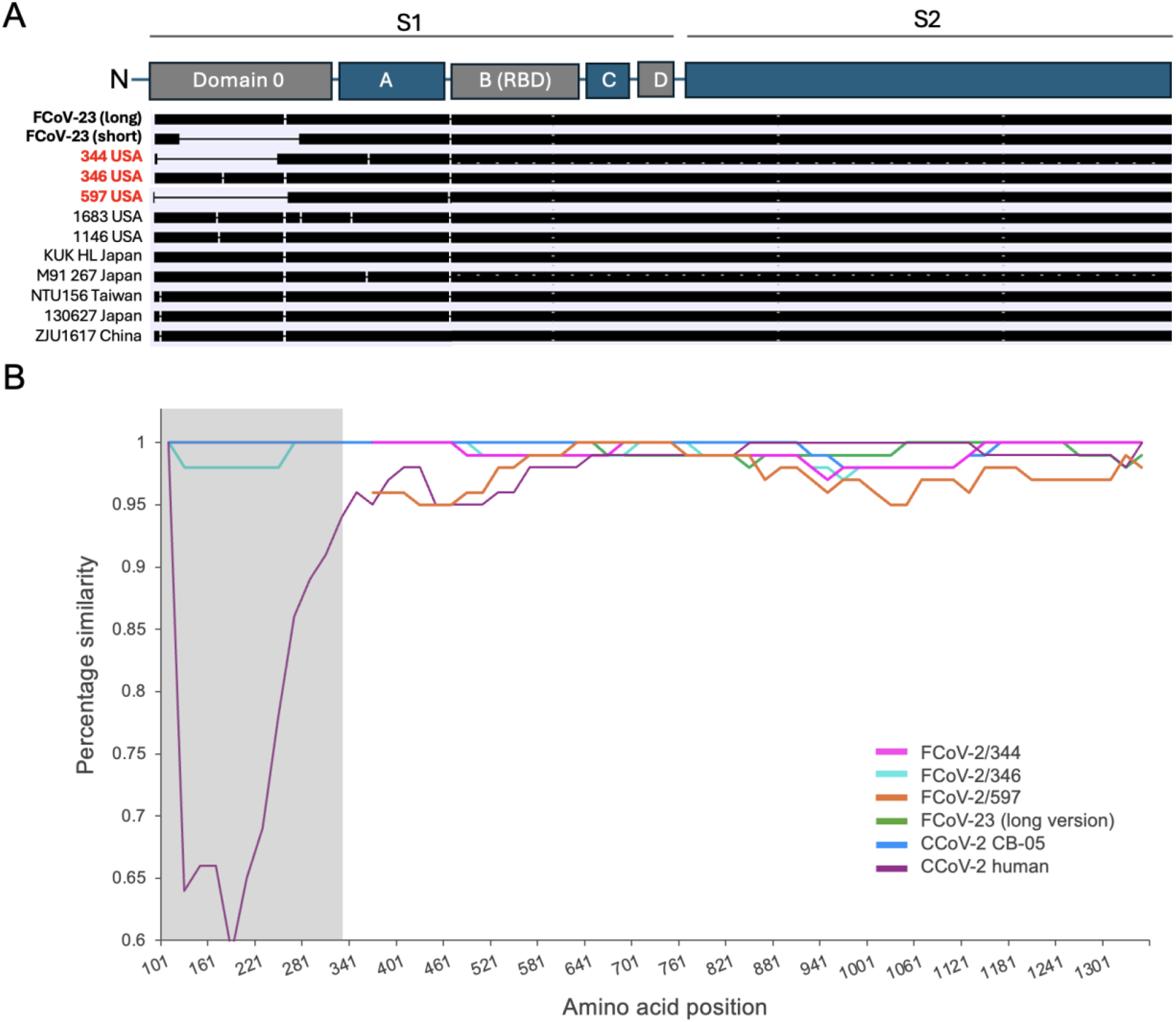
Genetic diversity of the S protein of FCoV-2, FCoV-23, and CCoV-2. On top is a graphical depiction of the different domains of the S protein. A. Graphical representation of the different sizes of the S protein from different FCoV-2 and FCoV-23 variants. Each horizontal black bar represents the length of the S protein of 12 FCoVs, including the three identified in this study (in red) and the long and short versions of FCoV-23 (in bold). Deletions are shown as thin horizontal lines and are all in domain 0. The short versions of FCoV-23, FCoV-2/344, and FCoV-2/597 have deletions of different sizes. FCoV-2/346 does not have a deletion in this region. RBD receptor binding domain. B. Similarity plot of the amino acid sequence of the S protein (1,341 amino acids). CCoV-2 NA09/2009/Greece/JF682842, which is the closest variant to FCoV-23, was used as a reference. Domain 0 is shaded in gray in accordance with the placement shown in the graphical depiction of S at the top of the figure. FCoV-2/344 and FCoV/346 feature deletions in this region; thus, comparisons in this area are not possible. The plot was generated using the Kimura 2-parameter distance model, with a 200 bp window and a 20 bp step. Sequences used in the plot: CCoV-2 CB-05: KP981644, CCoV-2 human: MZ420153, FCoV-23 (long version): PQ133182.

A similarity plot of the amino acid sequence of S, using the closest variant to FCoV-23 as a reference (CCoV-2 NA09/2009/Greece), showed that the FCoV-2/344, FCoV-2/346, FCoV-2/597, FCoV-23, and pCCoV-2 CB-05/2005/Italy exhibit over 90% amino acid similarity across the S protein sequence, even when including domain 0 for FCoV-2/346 and the long version of FCoV-23 (Figure 3). In contrast, domain 0 of the CCoV-2 detected in humans is highly dissimilar (Figure 3). Phylogenetic analysis of the amino acid sequence of S, excluding the deletion region, showed that the three FCoV-2 variants reported in this study form a distinct group, closely related to a group that includes FCoV-23 and other pCCoV-2 variants reported in Europe (Figure 4).

**Figure 4.**
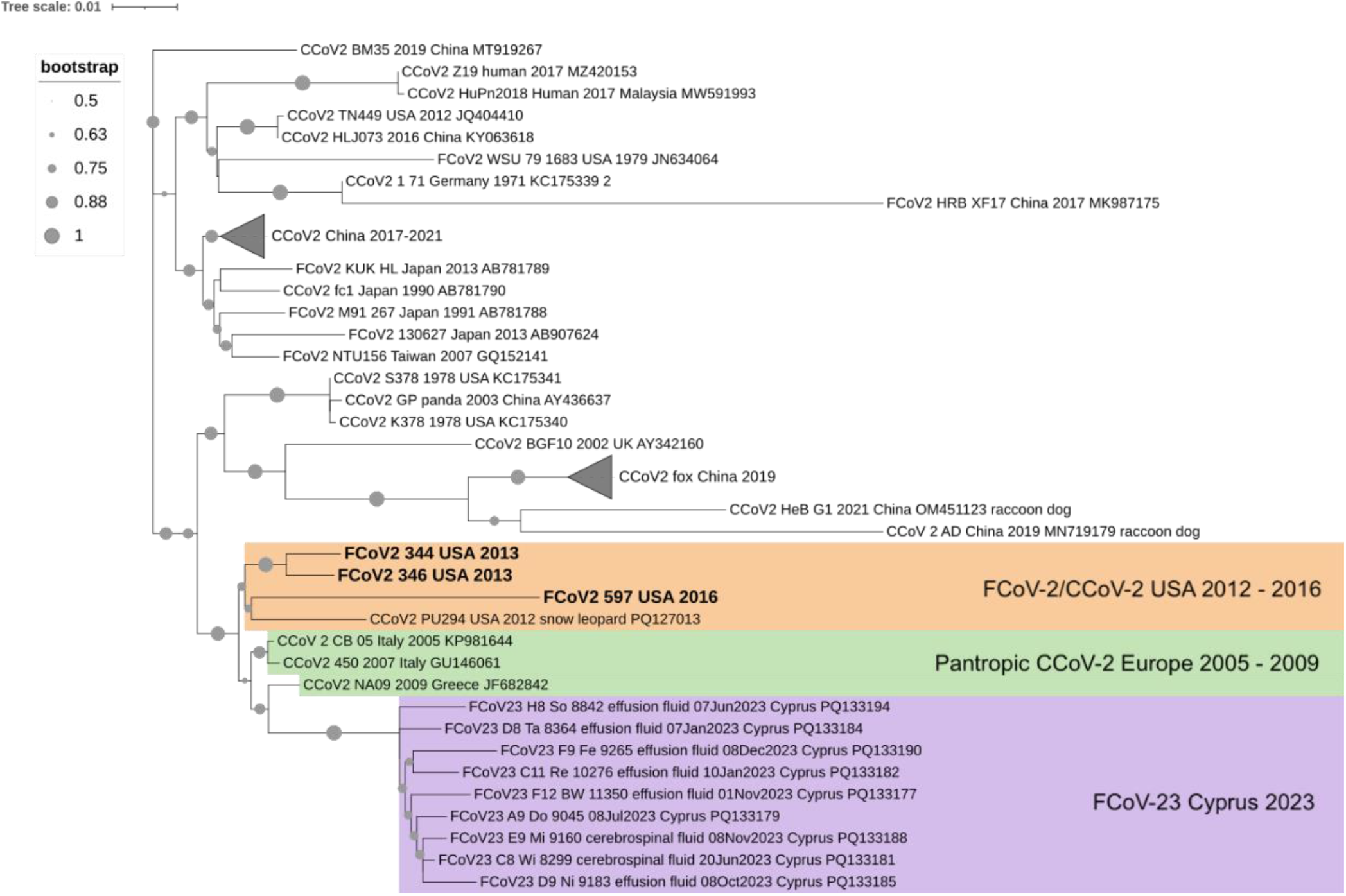
Maximum likelihood phylogenetic tree of the amino acid sequence of the S protein (1,081 amino acids) of FCoV and CCoV. The domain 0 region (where large deletions occur) was excluded. CCoV-2 and FCoV-2 detected in the USA between 2012 and 2016, including the three reported in this study (in bold), are in orange. Pantropic CCoVs reported in Europe are in green. FCoV-23 is in purple. Circles in the branches indicate bootstrap percentage values for 1,000 replicates. The size of each circle is proportional to the bootstrap value, and the scale is shown at the top left. Best-fitting nucleotide substitution model: GTR+G+I.

FCoV-2 RNA levels varied among the 20 samples tested from the three cats (Figure 5). For cat #597, FCoV-2 RNA levels ranged from 2.5 RNA copies/μl in feces to 93,174.5 RNA copies/μl in the mesenteric lymph (Appendix Table 4). For cat #344, the concentration of FCoV-2 RNA ranged from 5.3 RNA copies/μl in feces to 449,125 RNA copies/μl in the omentum. The quantification of FCoV-1 in the fecal sample from cat #344 revealed an RNA load of 22,543.5 RNA copies/μl (Appendix Table 4). For cat #346, FCoV-2 RNA levels were 47.7 RNA copies/μl in ascites and 434.5 RNA copies/μl in spleen (Appendix Table 4). The immunohistochemistry detection of FCoV-2 revealed extensive viral infection in the samples from cats #597 and #344 when compared to the spleen sample of cat #346 (Figure 6).

**Figure 5.**
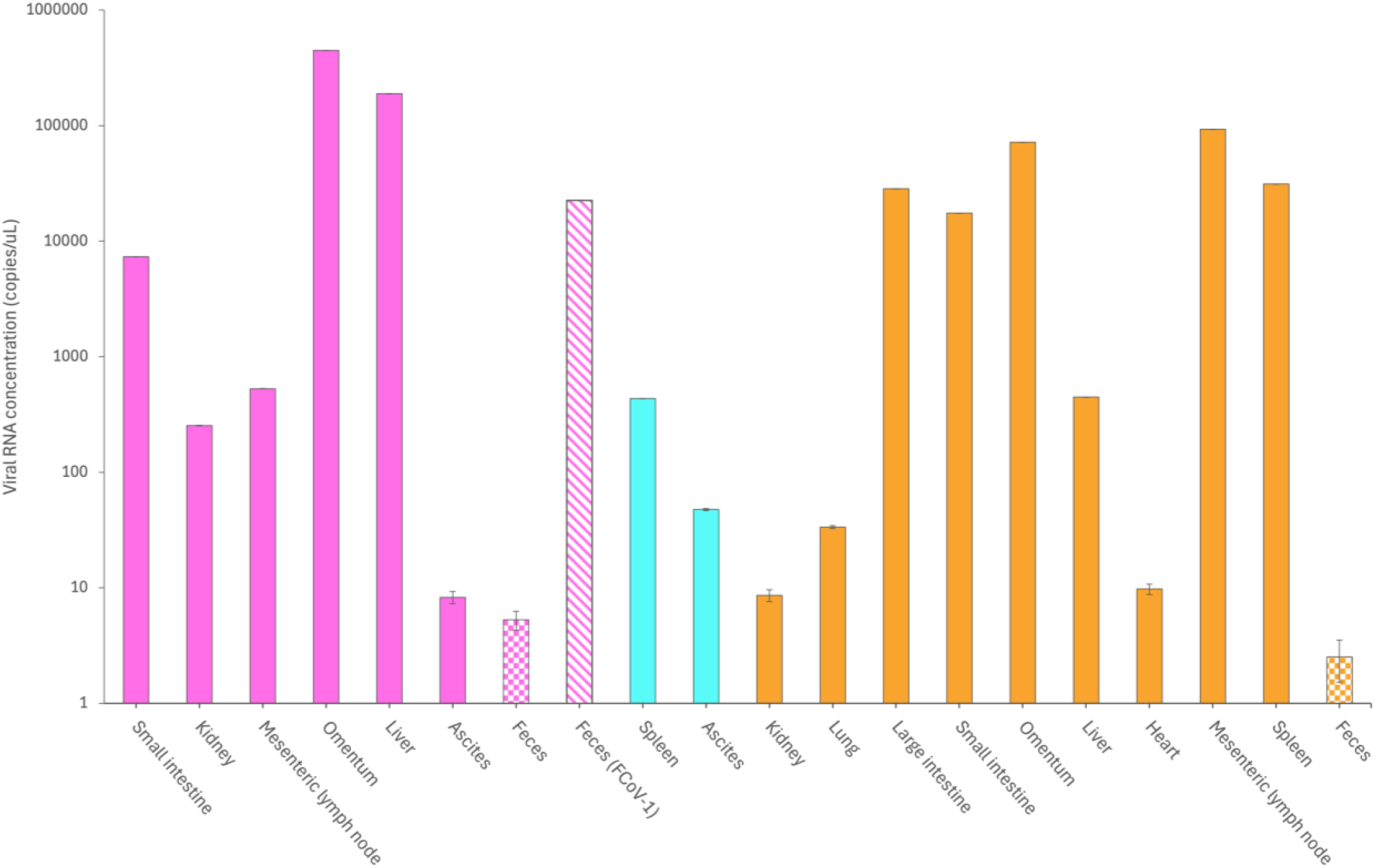
Quantification of FCoV-2 and FCoV-1 RNA in tissues, fluids, and feces of cats #344 (fuchsia), #346 (cyan), and #597 (orange). FCoV-1 RNA was detected and quantified only in the feces of cat #344 (bar with diagonal lines). Bars with squares represent the concentration of FCoV-2 RNA quantified in the feces of cats #344 and #597. The values shown are from two replicates.

**Figure 6.**
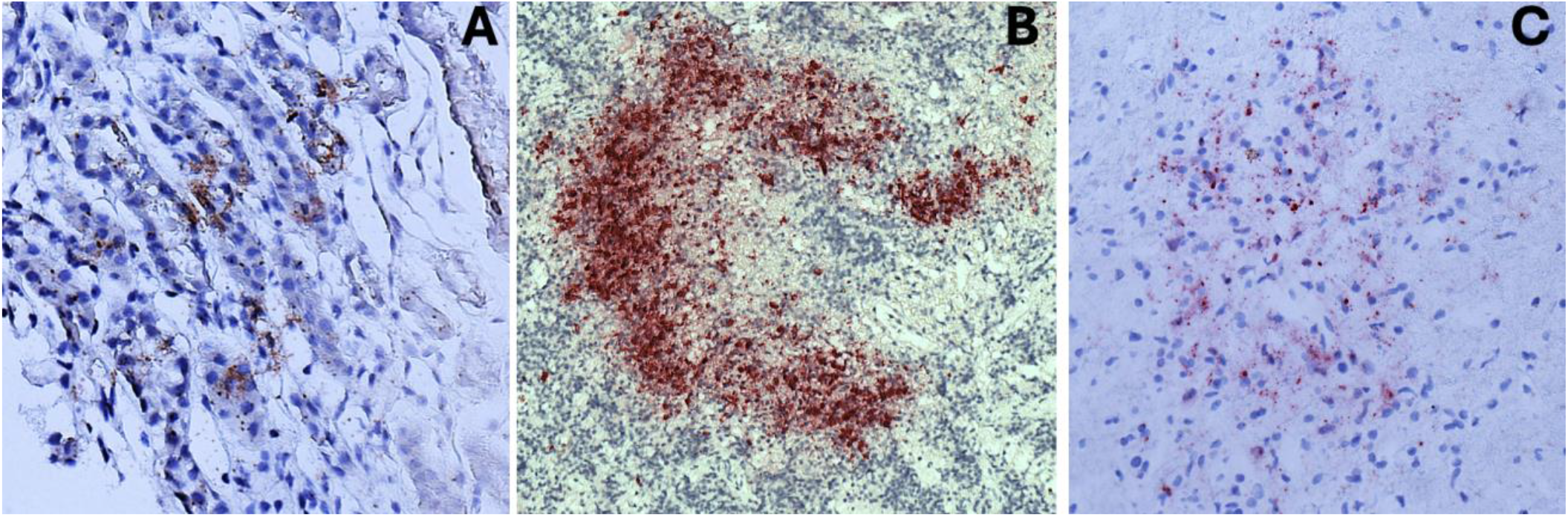
Detection of FCoV-2 (in burgundy) in tissues of cats #344, #597, and #346 by immunohistochemistry. A. Kidney of #344. B. Mesenteric lymph node of #597. C. Spleen of #346.

## Discussion

FCoV-23 is a highly pathogenic FCoV that emerged on the island of Cyprus, resulting in a widespread deadly outbreak among free-ranging cats (8). FCoV-23 resembles FCoV-2 as it is a recombinant of FCoV-1 and CCoV-2, which most likely exchanged the S gene with a pCCoV-2. The S protein of FCoV-23 is characterized by having two versions that differ in length. The short version is characterized by a variable-size deletion within domain 0 (Table 2). Variants displaying the deletion exhibit enhanced fusogenicity and cell entry, suggesting that the deletion is a significant pathogenicity factor for FCoV-23 (19). In this study, we conducted a molecular retrospective survey of FCoV-2 detected in three cats in the U.S. in 2013 and 2016, revealing two variants of FCoV-2 with different recombination breakpoints (Figure 1) and deletions in domain 0 of spike (Figure 3). Two of the three cats (#344 and #597) displayed signs for a prolonged time, including ascites (#344, #597) and upper respiratory signs and diarrhea (#597). We detected different deletions in domain 0 in the FCoV-2 detected in these two cats (Figure 3). The deletion size differed between individuals (153 amino acids in FCoV-2/344 and 212 amino acids in FCoV-2/597, Figure 3), but it was identical within each cat and was identified in all tissues and fluids examined (Table 1). The third cat (#346), daughter of #344, did not exhibit a history of persistent clinical signs, and the FCoV-2 detected in the two samples examined from this cat did not show deletions in S (Figure 3). Although the duration of infection of all the cats is unknown, we hypothesize that cats #344 and #597 had a prolonged infection because their medical history indicates that they were displaying clinical signs related to the infection with FCoV-2 (ascites, diarrhea) for at least two months. Persistent FCoV infection in cats has been widely reported (35, 36), and most cats with chronic diarrhea are infected with FCoV (37). Cat #346 was euthanized approximately one month after the initial development of ascites. Therefore, we assume that this cat was infected for a shorter duration than cat #344, which had ascites for nearly three months (from early March to late May 2013). Our results suggest that prolonged infection with FCoV-2 leads to deletions in domain 0 of S, and the viruses with the deletion rapidly spread within each host, causing extensive infection (Figure 6). Analogous to the ‘internal mutation’ hypothesis of FCoV-1, in which the accumulation of specific mutations leads to highly pathogenic FCoV-1 in persistently infected cats, we call this the ‘internal deletion’ hypothesis.

The transmission of FCoV-2 primarily occurs through the fecal-oral route, and the viral load in feces is crucial for viral transmission. For example, 80% (4 of 5) of cats experimentally inoculated with feces containing a low FCoV RNA concentration did not seroconvert or shed FCoV in their feces (38). In contrast, 100% (17 of 17) of those inoculated with feces containing a high viral RNA concentration rapidly seroconverted and began shedding the virus in their feces (38). Cats that are persistently infected and shed high titers of FCoV RNA in their feces are the most significant source of FCoV in multi-cat settings (35). Therefore, measuring viral RNA titer in feces can serve as a proxy for the likelihood of FCoV transmission. Cats #344 and #346 were living in the same house, and the FCoV-2 detected in these cats were 97.4% similar, had the same recombination breakpoints (Figure 1), and grouped together in the phylogenies of ORF1a (Figure 2) and S (Figure 4). The main difference between them was that FCoV-2/346 has the long version of S, while FCoV-2/344 has the short version of S (Figure 3). Although we detected FCoV-2 RNA with the short version in the feces of #344 and #597, the viral RNA load was very low (2.5 RNA copies/μl for FCoV-2/597 and 5.3 RNA copies/μl for FCoV-2/344, Appendix Table 4) when compared to the viral RNA load in the tissues and fluids (ranging from 828 ± 0.2 to 449,125.0 ± 1,438.9 RNA copies/μl for FCoV-2/344, and 9.8 ± 1.3 to 93,174.5 ± 1,167.4 RNA copies/μl for FCoV-2/597, Appendix Table 4) and with the RNA load of the FCoV-1 detected in the feces of #344 (22,543.5 ± 118.1, Figure 4). These results suggest that the FCoV-2 with the long version of the S gene was transmitted between cats #344 and #346, and the deletion observed in FCoV-2/344 developed within-host and spread systemically in cat #344 (Table 1). Supporting this and the ‘internal deletion’ hypothesis is the fact that 90% of the cats infected with FCoV-23 in Cyprus also exhibit deletions that vary in size and are specific to each individual (8). In the case of the Cyprus outbreak, high FCoV-23 RNA loads have been reported in feces; however, it remains unclear whether FCoV-23 displaying the long or short version of S is being shed (8). Overall, these results indicate that FCoV-2 with the long version of S was transmitted between cats #344 and #346, and within-host deletions in S emerged in the persistently infected cat (#344) and quickly spread systemically.

These results indicate that monitoring the evolution of the S gene during FCoV-2 outbreaks is important to determine if individuals with prolonged infections develop deletions in domain 0. For example, in an outbreak in Taiwan, the duration of FCoV-2 infection, measured from the onset of fever to death, ranged from 15 to 67 days in four cats (5). Although the S gene was sequenced in some of these cats, only the 3’-end was targeted; thus, the evolution of the 5’-end, where domain 0 is located, among cats with different infection times remains unknown. Phylogenetic and genetic comparisons of the entire S protein sequence revealed that the FCoV-2s reported in this study are closely related to pCCoV-2s and FCoV-23 (Figure 4). Additionally, their S protein is highly conserved, displaying over 90% amino acid similarity across the entire protein (Figure 3), including domain 0, which is highly divergent in other coronaviruses, such as CCoV-2 reported in humans (Figure 3). Considering the close genetic relationship with pCCoV-2 and FCoV-23, and the detection of highly pathogenic variants with domain 0 deletions in the U.S., it’s crucial to start surveillance programs for FCoV in multi-house settings like shelters, as well as free-ranging domestic cats in the U.S., to prevent a large-scale outbreak similar to the one in Cyprus.

Our results suggest that rapid molecular detection and characterization of FCoV are essential for preventing the development and spread of highly pathogenic viruses that exhibit deletions in the S gene. In the feces of cat #344 we detected co-infection with both FCoV-1 and FCoV-2. The viral RNA load of FCoV-1/344 in feces was higher than FCoV-2/344 (Appendix Table 2), but this FCoV-1 lacked any of the pathogenicity markers in S (i.e. no mutations were identified in the S1/S2 cleavage site or in residue ‘1058’). In this case, the pathogenic variant that spread systemically and caused FIP was a FCoV-2 displaying a deletion in S. Therefore, detection methods targeting only the S of FCoV-1 can detect FCoV-1 without markers of pathogenicity while overlooking co-infection with highly pathogenic FCoV-2 variants. Likewise, detection methods that focus on regions outside the S gene, which are more conserved, would have also missed the co-infection, as FCoV-2/344 and FCoV-1/344 were highly similar (94.1% pairwise similarity, excluding the recombinant region). In this study, we used hybridization capture targeting FCoV-1, FCoV-2, CCoV-1, and CCoV-2. This method allowed us to detect both co-infection between FCoV-1 and FCoV-2, as well as variants of FCoV-2 with major deletions not only in S, but also in other genes (Table 2). Given the complexity of diagnosing FIP in cats, it is emphasized that advanced techniques capable of detecting deletions and simultaneously differentiating between genotypes, like the one used in this study, are essential.

In conclusion, we propose an internal deletion model for within-host generation of highly pathogenic FCoV-2 through the loss of spike domain 0 (Figure 3). We further propose that FCoV-23 is not in itself a novel FCoV-2 variant, but is a highly impactful presentation of this mechanism based on the high population density of free-roaming cats on the island of Cyprus.

## Supporting information

Appendix inforamtion

## Acknowledgments

We thank Dr. Mandi de Mestre for allowing us to use the ddPCR system to quantify viral RNA and Donald Miller for teaching us how to use it. This research was funded by the Morris Animal Foundation (Grant ID D25FE-713: FCoV-23 Genomic Surveillance Core) and Cornell Feline Health Center. Cornell University’s Institutional Animal Care and Use Committee reviewed and approved the study procedures (approval number 2012-0116).

## Author’s contributions

XAO-C conceptualized the study, designed the experiments, performed data analysis, and wrote the initial draft of the manuscript. AS performed experiments, collected data, and revised the manuscript. KS performed experiments and revised the manuscript. LBG contributed to the analysis and interpretation of data, revised the manuscript, and acquired the funding for the study. GBW contributed to the literature review, revised the manuscript, and acquired the funding for the study.

