## Appendix inforamtion for "Identification of within-host deletions in domain 0 of the spike gene of highly pathogenic feline coronavirus type 2 from the United States"

### Table of contents


| | |
|---|---|
| **Appendix Table 1.** Size of S, in nucleotides (nt) and amino acids (AA) for available FCoV-2 variants. | **Page 2** |
| **Appendix Table 2.** List of the sequences used in the different analyses and phylogenies of this study. For all variants included, the sequence name includes the type of CoV (FCoV-1, -2, CCoV-1, -2), year and place of collection, and accession number. | **Page 3** |
| **Appendix Table 3.** Primers and probes used for quantifying the RNA of FCoV-1 and FCoV-2. | **Page 7** |
| **Appendix Table 4.** Viral RNA quantification in the samples collected from three cats (#344, #346, #597). | **Page 8** |

**Appendix Table 1.** Size of S, in nucleotides (nt) and amino acids (AA) for available FCoV-2 variants.

| **Variant name (accession number)** | **Collection place, date** | **Collection site** | **Length of S (nt/AA)** | **Reference** |
|---|---|---|---|---|
| FCoV-2 1146 (AY994055) | USA ,1979 | Lung, liver, spleen | 4356/1452 | (1) |
| FCoV-2 1683 (JN634064) | USA, 1979 | Mesenteric lymph node and intestinal wash | 4350/1450 | (1) |
| FCoV-2 KUK-H/L (AB781789) | Japan, 1987 | Spleen | 4362/1454 | (2) |
| FCoV-2 M91-267 (AB781788) | Japan, 1991 | Spleen | 4359/1453 | (2) |
| FCoV-2 NTU156 (GQ152141) | Taiwan, 2007 | Pleural effusion | 4359/1453 | (3) |
| FCoV-2 130627 (AB907624) | Japan, 2013 | Ascites | 4359/1453 | (4) |
| FCoV-2 HRB-17 (MK987175) | China, 2017 | Blood | 4359/1453 | (5) |

**Appendix Table 2.** List of the sequences used in the different analyses and phylogenies of this study. For all variants included, the sequence name includes the type of CoV (FCoV-1, -2, CCoV-1, -2), year and place of collection, and accession number.

| Sequence ID | Used in mapping process | Used in recombination analysis | Used in phylogeny of ORF1a | Used in phylogeny of S |
|---|---|---|---|---|
| CCoV1_23_Italy_2003_KP849472 | yes | yes | no | no |
| CCoV2_1_71_Germany_1971_KC175339 | yes | yes | no | yes |
| CCoV2_15_2020_UK_MT906864 | yes | yes | no | no |
| CCoV2_191101SY_China_2019_OQ540910 | yes | yes | no | yes |
| CCoV2_191110SY_China_2019_OQ540919 | yes | yes | no | yes |
| CCoV2_191133SY_China_2019_OQ540914 | yes | yes | no | yes |
| CCoV2_191140SY_China_2019_OQ540915 | yes | yes | no | yes |
| CCoV2_450_2007_Italy_GU146061 | no | no | no | yes |
| CCoV2_7_2020_UK_MT906865 | yes | yes | no | no |
| CCoV2_AD_China_2019_MN719179 | yes | yes | no | no |
| CCoV2_B135_JS_2018_China_MT114544 | yes | yes | no | yes |
| CCoV2_B194_GZ_2019_China_MT114543 | yes | yes | no | yes |
| CCoV2_B203_GZ_2019_China_MT114542 | yes | yes | no | yes |
| CCoV2_B363_ZJ_2019_China_MT114541 | yes | yes | no | yes |
| CCoV2_B447_ZJ_2019_China_MT114540 | yes | yes | no | yes |
| CCoV2_B600_ZJ_2019_China_MT114539 | yes | yes | no | yes |
| CCoV2_B639_ZJ_2019_China_MT114538 | yes | yes | no | yes |
| CCoV2_BGF10_2002_UK_AY342160 | no | no | no | yes |
| CCoV2_BM35_2019_China_MT919267 | no | no | no | yes |
| CCoV2_CB_05_Italy_2005_KP981644 | yes | yes | no | yes |
| CCoV2_fc1_Japan_1990_AB781790 | no | no | no | yes |
| CCoV2_GDX9_China_2020_MZ320954_ | yes | yes | no | yes |
| CCoV2_GH4_2_2020_China_OM950729 | yes | yes | no | yes |
| CCoV2_GH8_2_2020_China_OM950728 | yes | yes | no | yes |
| CCoV2_GP_2003_China_AY436637_giant_panda | no | no | no | yes |
| CCoV2_HeB_G1_2021_China_OM451123 | yes | yes | no | yes |
| CCoV2_HLJ071_2016_China_KY063616 | yes | yes | no | yes |
| CCoV2_HLJ072_2016_China_KY063617 | yes | yes | no | yes |
| CCoV2_HuPn2018_2017_Malaysia_MW591993_human | yes | yes | no | yes |
| CCoV2_K378_1978_USA_KC175340 | yes | yes | no | yes |
| CCoV2_NA09_2009_Greece_JF682842 | no | no | no | yes |
| CCoV2_PK02_2019_Brazil_OP179857 | yes | yes | yes | no |

| | | | | |
|---|---|---|---|---|
| CCoV2_PU294_2012_USA_snow_leopard_PQ127013 | yes | yes | yes | yes |
| CCoV2_S378_1978_USA_KC175341 | yes | yes | no | yes |
| CCoV2_SD_f3_2021_China_OM451122 | yes | yes | no | yes |
| CCoV2_SMU_8_2020_China_ON107244 | yes | yes | no | yes |
| CCoV2_Z19_Haiti_2017_MZ420153_human | yes | yes | no | yes |
| CCoV2a_HLJ073_2016_China_KY063618 | yes | yes | no | yes |
| CCoV2a_TN449_USA_2012_JQ404410 | yes | yes | no | yes |
| CCoV2b_NTU336_2008_Taiwan_GQ477367 | yes | yes | no | no |
| CCoV2c_A76_1976_USA_JN856008 | yes | yes | no | no |
| FCoV1_11HRBef_China_2016_KY566210 | yes | yes | no | no |
| FCoV1_13HRBef_China_2016_KY566211 | yes | yes | yes | no |
| FCoV1_1HRBef_2016_KY292377 | yes | yes | yes | no |
| FCoV1_26M_UK_2013_KP143512 | yes | yes | no | no |
| FCoV1_27C_UK_2013_KP143507 | yes | yes | yes | no |
| FCoV1_65F_UK_2013_KP143509 | yes | yes | no | no |
| FCoV1_67F_UK_2013_KP143510 | yes | yes | no | no |
| FCoV1_80F_UK_2013_KP143511 | yes | yes | no | no |
| FCoV1_Black_EU186072 | yes | yes | yes | no |
| FCoV1_C1Je_UK_2002_DQ848678 | yes | yes | yes | no |
| FCoV1_DCB091_USA_2008_PP854702 | yes | yes | yes | no |
| FCoV1_felix_Germany_2012_MG893511 | yes | yes | yes | no |
| FCoV1_FM3_USA_2020_sand_cat_OP542206 | yes | yes | yes | no |
| FCoV1_HLJ_HRB_China_2016_KY566209 | yes | yes | no | no |
| FCoV1_OM1164_pallas_cat_USA_2008_PP854701 | yes | yes | yes | no |
| FCoV1_RM_USA_2002_FJ938051 | yes | yes | yes | no |
| FCoV1_SB22_Brazil_2016_MH817484 | yes | yes | yes | no |
| FCoV1_UU10_Netherlands_2007_FJ938059 | yes | yes | no | no |
| FCoV1_UU11_Netherlands_2007_FJ938052 | yes | yes | no | no |
| FCoV1_UU15_Netherlands_2007_FJ938057 | yes | yes | yes | no |
| FCoV1_UU17_Netherlands_2007_HQ012367 | yes | yes | yes | no |
| FCoV1_UU2_USA_1993_FJ938060 | yes | yes | yes | no |
| FCoV1_UU20_Netherlands_2007_HQ392471 | yes | yes | yes | no |
| FCoV1_UU22_Netherlands_2007_GU553361 | yes | yes | no | no |
| FCoV1_UU23_Netherlands_2007_GU553362 | yes | yes | no | no |
| FCoV1_UU24_Netherlands_2008_HQ012370 | yes | yes | yes | no |
| FCoV1_UU30_Netherlands_2008_HQ392472 | yes | yes | yes | no |
| FCoV1_UU31_Netherlands_2008_HQ012371 | yes | yes | yes | no |
| FCoV1_UU4_Netherlands_2007_FJ938054 | yes | yes | yes | no |
| FCoV1_UU5_Netherlands_2007_FJ938056 | yes | yes | yes | no |

| | | | | |
|---|---|---|---|---|
| FCoV1_UU7_Netherlands_2007_FJ938053 | yes | yes | yes | no |
| FCoV1_UU8_Netherlands_2007_FJ938055 | yes | yes | no | no |
| FCoV1_UU9_Netherlands_2007_FJ938062 | yes | yes | yes | no |
| FCoV2_130627_Japan_2013_AB907624 | no | no | no | yes |
| FCoV2_79_1146_USA_1979_AY994055 | yes | yes | yes | no |
| FCoV2_HRB_XF17_China_2017_MK987175 | no | no | no | yes |
| FCoV2_KUK_HL_Japan_2012_AB781789 | no | no | no | yes |
| FCoV2_M91_267_Japan_2012_AB781788 | no | no | no | yes |
| FCoV2_NTU156_Taiwan_2007_GQ152141 | yes | yes | yes | yes |
| FCoV2_WSU_79_1683_USA_1979_JN634064 | yes | yes | yes | yes |
| FCoV23_A9_Do_9045_08Jul2023_Cyprus_PQ133179 | yes | yes | yes | no |
| FCoV23_C11_Re_10276_2023_Cyprus_PQ133182 | yes | yes | yes | no |
| FCoV23_C8_Wi_8299_2023_Cyprus_PQ133181 | yes | yes | yes | no |
| FCoV23_D8_Ta_8364_2023_Cyprus_PQ133184 | yes | yes | yes | no |
| FCoV23_D9_Ni_9183_2023_Cyprus_PQ133185 | yes | yes | yes | no |
| FCoV23_E9_Mi_9160_2023_Cyprus_PQ133188 | yes | yes | yes | no |
| FCoV23_F11_Gi_6590_2023_Cyprus_PQ133176 | yes | yes | yes | no |
| FCoV23_F12_BW_11350_2023_Cyprus_PQ133177 | yes | yes | yes | no |
| FCoV23_F9_Fe_9265_2023_Cyprus_PQ133190 | yes | yes | yes | no |
| FCoV23_H8_So_8842_2023_Cyprus_PQ133194 | yes | yes | yes | no |
| TGEV_138_2006_USA_KX900395 | yes | yes | no | no |
| TGEV_139_2006_USA_KX900396 | yes | yes | no | no |
| TGEV_140_2007_USA_KX900397 | yes | yes | no | no |
| TGEV_141_2007_USA_KX900398 | yes | yes | no | no |
| TGEV_142_2007_USA_KX900399 | yes | yes | no | no |
| TGEV_143_2008_USA_KX900400 | yes | yes | no | no |
| TGEV_144_208_USA_KX900401 | yes | yes | no | no |
| TGEV_145_2008_Mexico_KX900402 | yes | yes | no | no |
| TGEV_146_2008_USA_KX900403 | yes | yes | no | no |
| TGEV_147_2012_USA_KX900404 | yes | yes | no | no |
| TGEV_148_2013_USA_KX900405 | yes | yes | no | no |
| TGEV_149_2013_USA_KX900406 | yes | yes | no | no |
| TGEV_150_2013_USA_KX900407 | yes | yes | no | no |
| TGEV_151_2014_USA_KX900408 | yes | yes | no | no |
| TGEV_152_2014_USA_KX900409 | yes | yes | no | no |
| TGEV_153_2014_USA_KX900410 | yes | yes | no | no |
| TGEV_154_2014_USA_KX900411 | yes | yes | no | no |
| TGEV_AHHF_2015_China_KX499468 | yes | yes | no | no |
| TGEV_CH8438_2017_China_MW804449 | yes | yes | no | no |

| | | | | |
|---|---|---|---|---|
| TGEV_CHGX_2662_2019_China_MZ322950 | yes | yes | no | no |
| TGEV_CHN_SC_H_ON016092 | yes | yes | no | no |
| TGEV_HB_1988_USA_KX900394 | yes | yes | no | no |
| TGEV_HB1_2020_China_MZ368889 | yes | yes | no | no |
| TGEV_HE1_2015_China_KX083668 | yes | yes | no | no |
| TGEV_HLJ17_2017_China_MT522161 | yes | yes | no | no |
| TGEV_HN_2012_China_OP434397 | yes | yes | no | no |
| TGEV_HNSQ_2021_China_ON859974 | yes | yes | no | no |
| TGEV_HQ_2016_China_MT576083 | yes | yes | no | no |
| TGEV_HX_2012_China_KC962433 | yes | yes | no | no |
| TGEV_SCY_China_DQ443743 | yes | yes | no | no |
| TGEV_SHXB_2013_China_KP202848 | yes | yes | no | no |
| TGEV_TH_1998_China_KU729220 | yes | yes | no | no |
| TGEV_WH_1_2010_China_HQ462571 | yes | yes | no | no |
| TGEV_Z1986_2006_USA_KX900393 | yes | yes | no | no |

**Appendix Table 3.** Primers and probes used for quantifying the RNA of FCoV-1 and FCoV-2.

| Primer/ probe name | Sequence (5'- 3') | Target |
|---|---|---|
| 344_FCoV2_F | GGCTTAGGTACAGTCGATGAAG | S gene of FCoV-2 344/ 346/ 597 |
| 344_FCoV2_R | CATTAGCCACACCAGGTAACA | |
| 344_FCoV2_P | 5′-FAM-ACGTTGTACGGGTGGCTATGAYAT-BHQ-1 | |
| 344_FCoV1_F | CACCTCAGCTTGTCAAACAATAG | S gene of FCoV-1 344 |
| 344_FCoV1_R | GACTGCGATCTGACACTGTAAT | |
| 344_FCoV1_P | 5′-FAM-AATGCCCTTAATCTTGGTGCACGC-BHQ-1 | |

**Appendix Table 4.** Viral RNA quantification in the samples collected from three cats (#344, #346, #597).

| Cat ID | Sample ID/ type | Viral RNA quantification (RNA copies/µl)* | |
|---|---|---|---|
| | | FCoV-1 | FCoV-2 |
| 344 | 344.1 Small intestine | Neg | 7,335.5 ± 40.3 |
| | 344.2 Kidney | Neg | 253.5 ± 7.8 |
| | 344.3 Mesenteric lymph node | Neg | 528.5 ± 37.5 |
| | 344.4 Omentum | Neg | 449,125.0 ± 1,438.9 |
| | 344.5 Liver | Neg | 189,250.0 ± 6,717.5 |
| | 344.6 Feces | 22,543.5 ± 118.1 | 5.3 ± 1.4 |
| | 344.7 Spleen | Neg | n.o. |
| | 344.8 Ascites | Neg | 828 ± 0.2 |
| 346 | 346.1 Spleen | Neg | 434.5 ± 20.5 |
| | 346.2 Ascites | Neg | 47.7 ± 3.6 |
| 597 | 597.1 Kidney | Neg | 8.6 ± 0.1 |
| | 597.2 Lung | Neg | 33.5 ± 1.4 |
| | 597.3 Large intestine | Neg | 28,489.0 ± 42.4 |
| | 597.4 Small intestine | Neg | 17,534.0 ± 551.6 |
| | 597.5 Omentum | Neg | 71,820.0 ± 217.8 |
| | 597.6 Liver | Neg | 447.5 ± 13.4 |
| | 597.7 Heart | Neg | 9.8 ± 1.3 |
| | 597.8 Mesenteric lymph node | Neg | 93,174.5 ± 1,167.4 |
| | 597.9 Spleen | Neg | 31,273.0 ± 113.1 |
| | 597.10 Feces | Neg | 2.525 ± 0.7 |

*: values shown for two replicates; n.o.: not obtained
